# Multi-omic analysis of colorectal adenocarcinoma identifies a new subtype of myofibroblastic cancer-associated fibroblast expressing high level of B7-H3 and with poor prognosis value

**DOI:** 10.1101/2025.06.30.662277

**Authors:** Maelle Picard, Arnaud Guille, Pascal Finetti, Bernadette de Rauglaudre, Nadiya Belfil, Lenaïg Maescam, David Jeremy Birnbaum, François Bertucci, Emilie Mamessier

**Affiliations:** Aix Marseille Univ, INSERM U1068, CNRS, Institut Paoli-Calmettes, CRCM, “ Predictive Oncology laboratory”, Label “Ligue contre le cancer”, Marseilles, France; Department of Digestive Oncology and Gastro-Enterology, Timone Hospital, AP-HM, 13005 Marseille, France; Aix Marseille Univ, Department of Digestive Surgery, Hôpital Nord, Assistance Publique-Hôpitaux de Marseille, Marseille, France; Department of Biopathology, Paoli-Calmettes Institute, 13009 Marseilles, France; Department of Medical Oncology, Paoli-Calmettes Institute, 13009 Marseille, France

**Keywords:** B7-H3, cancer-associated fibroblast, colorectal cancer, oncogenesis, prognosis

## Abstract

The high mortality rate of colorectal cancer (CRC) combined with the lack of non-toxic and efficient personalized treatments makes it urgent to develop new targeted therapies for this disease. B7-H3 appears to be a good target as it is overexpressed in tumor tissue compared to normal tissue. However, B7-H3 is a molecule with ambivalent functions and is expressed by different cell types. This complexity contributed to the delay in identifying cell subtypes that express B7-H3 and their potential role in colorectal oncogenesis.

In this study, we used *in silico* bulk, single-cell, and spatial transcriptomic data to investigate the clinical and biological characteristics of tumors with high B7-H3 expression, the precise nature of cells expressing high level of B7-H3, and their temporal appearance during colorectal oncogenesis. We found that tumors with high B7-H3 expression corresponded to tumors with a predominant stroma composed mainly of fibroblasts. Among them, two subtypes of ecm-myCAF fibroblasts and pro-fibrotic pericytes specifically expressed high levels of B7-H3, the former being an independent factor for poor prognosis in CRC. Finally, by examining precancerous lesions, we report that fibroblast subtypes with high levels of B7-H3 appear early during oncogenesis, especially at the inflamed stage.

We also shed light on the fact that anti-B7-H3 immunotherapies might therefore preferentially target cells from the microenvironment rather than the tumor cells. This is particularly important to understand the mode of action of the anti-B7-H3 antibody-drug conjugate, which is currently being tested in clinical trials in several solid tumors.

**Highlights:**

- High level of *B7-H3* expression is associated with poor prognosis in colorectal cancer
- *B7-H3* is recurrently found expressed by two subtypes of cancer-associated fibroblasts (CAF): pro-fibrotic pericytes and ecm-myCAF
- *B7-H3*^high^ ecm-myCAF subtype is an independent poor prognosis for survival
- *B7-H3*^high^ ecm-myCAF is detectable in pre-cancerous inflamed colonic tissues

## Introduction

Colorectal cancer (CRC) is the third most common cancer in men and women and the second leading cause of cancer-related deaths worldwide, making it a major public health problem [1]. The 5-year overall survival rate is of 63%, all stages combined. This rate depends on the tumor stage at diagnosis, the presence of distant metastases and the possibility of surgical intervention [2]. Standard treatments for primary non-metastatic CRC consist in surgery and followed by systemic chemotherapy in the poor-prognosis cases. However, about 70% of patients will relapse and develop resistance to this treatment within 5 years, highlighting the low efficiency of current treatment on residual tumor cells. Currently, most patients with primary CRC remain without efficient personalized options, and the exploration of novel therapeutic approaches is crucial [3, 4].

Over the past decade, immunotherapy has gained significant attention due to its remarkable efficacy, including in advanced CRC patients [4]. Another promising approach for the development of effective and less toxic therapies is to target tumor cells *via* tumor cell surface proteins with antibody drug conjugates (ADCs) [5]. In numerous solid tumors, ADCs have been quickly overturning the therapeutic management [6]. Similarly, in CRC, the recent DESTINY CR02 trial revealed promising antitumor activity and favorable safety profile of trastuzumab deruxtecan in patients with HER2-positive metastatic tumors, including those with *RAS* mutations. The use of ADC in the non-metastatic setting has not been investigated so far, but remains an interesting option to better eradicate disseminated tumor cells and further control disease progression.

B7-H3, also known as CD276, has been described as an important member of the B7/CD28 family. B7-H3 is a type-1 transmembrane glycoprotein with extracellular immunoglobulin-like domains, the transcript of which is ubiquitously expressed in immune cells, and at low levels in normal tissues. B7-H3 has been found to be expressed by malignant tumor cells in a broad spectrum of human solid tumors [7–12]. Overexpression of its transcripts from bulk tissue often correlated with poor-prognosis factors and poor clinical outcome in the majority of patients [7, 8, 10, 13]. More precisely, B7-H3 was shown to be associated with poor biological features, such as stemness [14], tumor angiogenesis [13], epithelial-to-mesenchymal transition [15, 16], and inflammation [17] and is closely related to tumor progression and survival [18] in CRC.

This led to a broad consensus on targeting B7-H3, as more than 30 clinical trials are currently registered on clinicaltrials.gov, suggesting that B7-H3 is a good target in the field of antibody-based therapeutics.

However, in CRC, the expression of B7-H3 has also been reported in the tumor-associated vasculature and in the general fibroblast population [19]. Cancer-associated fibroblasts (CAFs) are the most abundant stromal cells in CRC tumors and play a central role by influencing cancer cells proliferation, invasion, and metastasis, as well as angiogenesis and immune response by remodeling the extracellular matrix (ECM), secreting soluble factors (chemokines and growth factors), and modulating the microbial community [20, 21]. These functions are performed by different subtypes of fibroblasts. The prevailing classification of CAFs includes three main subtypes: myofibroblastic CAF (myCAF), inflammatory CAF (iCAF), and antigen-presenting CAF (apCAF), which have been described primarily in pancreatic and breast cancers, although other subtypes have been also found selectively [22, 23]. For now, it is not known if all CAFs or only subsets of CAFs express B7-H3. Given their involvement in most of the key steps of tumorigenesis, targeting CAF appears to be an attractive therapeutic option. However, targeting the wrong CAF subtypes can have dramatic consequences. Indeed, in pancreatic cancer, preclinical trials targeting the prominent stroma have been unsuccessful: non-specific depletion of fibroblasts leads to an increased proportion of the poorly differentiated, more aggressive subtype of pancreatic adenocarcinoma [24].

In this study, we wanted to clarify the expression of B7-H3 in CRC tumors, both as a biomarker and therapeutic target, to further our understanding on the potential mode of action of ongoing anti-B7-H3 immunotherapies. Notably, we investigated the clinical and biological characteristics of tumors enriched in *B7-H3*, the precise nature of cells expressing high level of B7-H3, and their temporal appearance during colorectal oncogenesis.

## Materials and Methods

### Datasets and Samples

#### Bulk transcriptome datasets

We retrospectively collected clinicopathological and gene expression data of CRC samples from 22 public data sets (Supplementary Table 1). We obtained 3964 samples profiled using DNA microarrays or RNA-seq at different stage of oncogenesis: 182 for normal colon, 222 for inflammatory bowel disease, 18 for polyps, 3,333 for primary tumors, and 209 for metastatic samples.

We also used cancer cell lines RNA sequencing data from the Dependency Map (DepMap) portal (https://depmap.org/portal) for analysis of RNA-protein correlation of B7-H3 expression.

#### Single-cell transcriptome datasets

Single-cell RNA sequencing (scRNAseq) data were obtained from publicly available datasets (CRC; GSE132465 and GSE144735 [25], mCRC; GSE178318 [26] and Atlas; Chu Xiaojing *et al.* [27]. Each dataset was analyzed independently to bypass batch effects, with same processing as authors. ScRNA-seq counts data under accession codes GSE132465 and GSE144735 were downloaded and processed using Seurat (v4.3). This dataset includes 29 primary CRC samples and 16 matched normal mucosa samples from 29 patients. GSE178318 includes 6 metastatic CRC samples from 6 patients. Processed data of single cell human colorectal atlas were downloaded from https://www-nature-com.proxy.insermbiblio.inist.fr/articles/s43018-024-00807-z#data-availability. This dataset includes data on tumor tissues from 124 patients, para-cancerous tissues from 78 patients, polyps from 9 patients, inflamed tissues from 23 patients, uninflamed tissues from 11 patients and healthy tissues from 36 patients.

#### Formalin-fixed paraffin-embedded samples

For immunofluorescence, we used the prospective cohort CTC-Colon cohort (CTC-Côlon – IPC 2015-020, NCT03256084) promoted by the Institut Paoli-Calmettes (Marseille, France). It includes patients with a pathological diagnosis of primary colic adenocarcinoma, confirmed by an expert pathologist of the Paoli-Calmettes Institute (Marseille, France). Main inclusion criteria were: patient of more than 18 years old; pathological diagnosis of colic adenocarcinoma, metastatic or not; before any treatment with systemic chemotherapy; signed informed patient’s consent; available samples formalin-fixed paraffin-embedded (FFPE). Surgical specimens were collected at the pathology department of the Paoli-Calmettes Institute, processed, and fixed in buffered formalin, paraffin-embedded per the standard-of-care for clinical pathology procedures. FFPE samples were obtained from pathology department archives following a Clinical Outcomes study (COS) authorization. Sample preparation, staining and read out were performed by members of the Predictive Oncology (PO) department (CRCM/IPC headed by Pr. Bertucci and Dr. Mamessier).

### Bulk transcriptome analysis

#### Gene expression

Before analysis, the pre-processing of data included two successive steps. The first one was to normalize each data set separately: we applied quantile normalization for the available processed data from non-Affymetrix-based sets, and Robust Multichip Average (RMA) with the non-parametric quantile algorithm for the raw Affymetrix-based data sets. Normalization was done in R using Bioconductor and associated packages [28]. In a second step, hybridization probes were mapped across the different technological platforms represented as previously reported [29]. When multiple probes mapped to the same GeneID, we retained the one with the highest variance in a particular data set.

Next, we corrected the 22 studies for batch effects using z-score normalization [30]. Briefly, for each *B7-H3* expression value in each study separately, the value was transformed by subtracting the mean of the gene in that dataset divided by its standard deviation in the primary CRC samples. Analysis was done by using binary values using the quartile of *B7-H3* expression level in the primary samples. B7-H3^Low^ was defined as the first quartile and B7-H3^High^ included the second through forth quartiles. Because B7-H3 is a family member of immune checkpoint B7/CD28 family, we searched for correlations of its expression in tumours with several molecular classification and immune variables. We thus applied several molecular or immune multigene classifiers to each tumour in each data set separately: Consensus molecular subtype (CMS) of CRC [31], CRCassigner-30 gene signatures [32], CAF signatures [33–35], ESTIMATE scores (Immune infiltration, Stromal infiltration, Tumor purity) [36], ICR STS signatures of 24 different innate and adaptative immune cell subpopulations defined by Bindea *et al.* [37], the tertiary lymphoid structures (TLS) signature [38], T-cell-inflamed signature [39], the cytolytic activity score [40], the pathway activation score of TGFβ [41], and the antigen processing machinery signature (APMS) score [42].

To decipher the biological pathways associated with *B7-*H3 expression in primary CRC samples, we applied a supervised analysis to expression profiles of the 459 TCGA samples (learning set) to search for genes differentially expressed between the “*B7-H3*-high” versus “*B7-H3*-low” classes. We used a moderated t-test with empirical Bayes statistic included in the limma R packages. False discovery rate [43] was applied to correct the multiple testing-hypothesis: the significant genes were defined by p<5%, q<5%, and fold change (FC) superior to |1.5x|. The robustness of the resulting gene list was tested in the validation set of 2,874 remaining samples (663 “*B7-H3*-low” samples and 2211 “*B7-H3*-high” samples) by computing for each tumor a metagene-based prediction score defined by the difference between the “metagene *B7-H3*-up” (mean expression of all genes upregulated in the “*B7-H3*-high” class) and the “metagene *B7-H3*-low” (mean expression of all genes upregulated in the “*B7-H3*-low” class). This score was then compared between the “*B7-H3*-high” and “*B7-H3*-low” samples. Ontology analysis of the resulting gene list was based on the Database for Annotation, Visualization and Integrated Discovery (DAVID; http://david.abcc.ncifcrf.gov/) and the GSEA processes.

#### Statistical analysis

The continuous variables are presented using median and range, whereas the discrete values are presented using number and percentage. The correlations between *B7-H3* expression-based groups and clinicopathological variables were calculated with the Student’s t-test or Fisher’s exact test when appropriate. The clinical endpoint of our prognostic analysis was the disease-free survival (DFS), calculated from the date of diagnosis until the date of disease relapse or death, the metastasis-free survival (MFS), calculated from the date of diagnosis until the date of first distant metastasis, and overall survival (OS), calculated from the date of diagnosis until the date of death. The follow-up was measured from the date of diagnosis to the date of last news for event-free patients. Univariate and multivariate prognostic analyses for DFS were done using Cox regression analysis (Wald test). The variables tested in univariate analysis included the sample groups based on *B7-H3* expression (“B7-H3-low”, “B7-H3-high”), patients’ age and sex (male, female), pTNM status, grade of differentiation, mismatch repair status (MMR), and the CMS classification. Multivariate analysis incorporated all variables with a p-value inferior to 5% in univariate analysis. Survival was calculated using the Kaplan-Meier method and curves were compared with the log-rank test. The correlations of molecular variables with “B7-H3-high” *versus* “B7-H3-low” groups were assessed using a logistic regression analysis using the glm function (significance estimated by specifying a binomial family for model with a logit link). All statistical tests were two-sided at the 5% level of significance. Survival analysis was done using the survival package (version 2.43) in the R software (version 3.5.2, https://cran.r-project.org).

### Single-cell statistical analysis

Briefly, for all datasets, the counts data were log-normalized, and variance stabilizing transformation (VST) method was used to identify the top 2000 highly variable features before performing Principal Component Analysis (PCA) with 50 components. Then, Harmony was employed to remove batch effects, using patient as a covariate and setting the number of dimensions to 40. To identify the main cell clusters, k-nearest neighbors (k-NN) and Louvain algorithm with resolution set at 0.1 were used. Author’s cells annotations were then used to identify fibroblast/stromal cells. For GSE178341, cell’s annotation was done with symphony using GSE178341 as reference. Finally, single-cell genes signatures scoring was done with the UCell package [44].

### Spatial transcriptomic

Spatial transcriptomic dataset was recovered from (https://www.10xgenomics.com/products/visium-hd-spatial-gene-expression/dataset-human-crc; doi: https://doi.org/10.1101/2024.06.04.597233). This dataset includes 5 patients, including 3 CRC samples and 2 normal colon tissues, analysed by spatial transcriptomic Visium HD platform (10X Genomics). B7-H3 level of expression was evaluated using LoopBrowser 8.

### Immunofluorescence

#### Region of Interest selection

Tissue sections of 3 µm were cut with a microtome (Epredia™ HM 340E Electronic Rotary Microtome, Fisher Scientific, US) and mounted on yellow slides Klinipath™ (Avantor, US). Slides were dried at room temperature before being coloured by Hematoxylin-Eosin-Safran. Slides were scanned using the NanoZoomer 2.0-HT™ (Hamamatsu Photonics, Japan). Region of interest (ROI) including healthy colon tissue, CRC or a mix of both were identified using the CaloPix software (version 2.10.16).

#### Staining

We selected 5 patients with a locally advanced CRC FFPE sample from our CTC-Colon cohort, presenting both healthy and malignant regions. Serial 3µm tissue sections were cut with a microtome (Epredia™ HM 340E Electronic Rotary Microtome, Fisher Scientific, US) and mounted on TOMO® Adhesion Microscope Slides (Matsunami GlassSlides, Japan), dried at room temperature before being incubated overnight at 56°C for a first step of dewaxing. Slides underwent a 10-minute-long wash of Histolemon® (CARLO ERBA, Italia) to complete dewaxing, then a progressive ethanol bath to rehydrate the tissue. Slides underwent heat-induced epitope retrieval in Tris-EDTA buffer (pH=9.0) following PT Link procedure (Dako Agilent, US).

To quench FFPE-tissue natural autofluorescence, slides were incubated in a solution of Sudan Black B 0.3% (SBB, Sigma Aldrich, US) ethanol 70% for 15 minutes. Slides were washed with Tween20 (Euromedex, France) 0.02% PBS (Gibco, Thermo Fischer Scientific, US) 1X for 15 minutes then PBS 1X for 5 minutes. Tissues were permeabilized with Tritonx100 (Euromedex, France) 0.5% PBS 1X for 15 minutes and saturated with Normal Horse Serum (NHS) 5% PBS 1X for 30 minutes.

An antibody panel targeting B7-H3 (*catalogue number #66481-1-Ig, clone 1E7D1, uncoupled, diluted to 1:25, Proteintech plus secondary antibody anti-Mouse Cy3 Jackson*) in tumor cells (*pan-KRT, catalogue number #53-9003-82, clone AE1/AE3, primary coupled with Alexa fluor 488, diluted to 1:50, eBioscience*) and in CAF (*VIM, catalogue number #130-127-018, clone REAL1008, primary coupled with APC, diluted to 1:50, Miltenyi*) was built. Antibodies mix was incubated 1 hour at room temperature and obscurity. Slides were washed several times by serial baths with saturation solution, then PBS 1X and finally with miliQ water before being slipped with Prolong antifade mounting medium (ThermoFisher Scientific, US).

#### Image Acquisition

ROI were imaged with Confocal Microscope (LSM800, Carl Zeiss, Germany) at X25 immersion magnification using appropriate exposure times and 2×2 tile scan. ROI were acquired and analysed with automatized software (Zen Black, version 3.3.89.0000, and ZEN 3.3 (blue edition), Carl Zeiss, Germany). Complementary acquisitions of the whole tissue slides were performed using a fluorescent scanner (Nanozoomer s60 v2 Digital slide scanner, Hamamatsu, Japan) to localized ROI within the tissue to guarantee the representativity of acquisitions.

## Results

### B7-H3 expression level is associated with clinicopathological parameters of primary CRC

Overexpression of *B7-H3* was previously reported in a series of 316 CRC cases [19]. To confirm this finding in a larger series of CRC, we collected a total of 3,333 CRC samples. In this series, the median age of patients was 69 years (range, 22-97 years), 52% of the patients were men and the tumor location was from the left colon in 52% of cases. Analyzed tumors were more frequently at a localized stage (stage 1 – 2, 63% of cases), pathological (p)T3 stage (76% of cases), pN0 stage (70% of cases were pN0), and pM0 stage (90% of M0), with a majority of grade 2 (78% of cases) and MSS tumors (82% of cases) (Table 1). We examined the level of *B7-H3* expression in our series compared to normal tissue and found that *B7-H3* was consistently overexpressed in CRC (*P*=4.67 x 10^-42^, Figure 1A). This confirmed the interest to explore further *B7-H3* expression in CRC.

**Figure 1.**
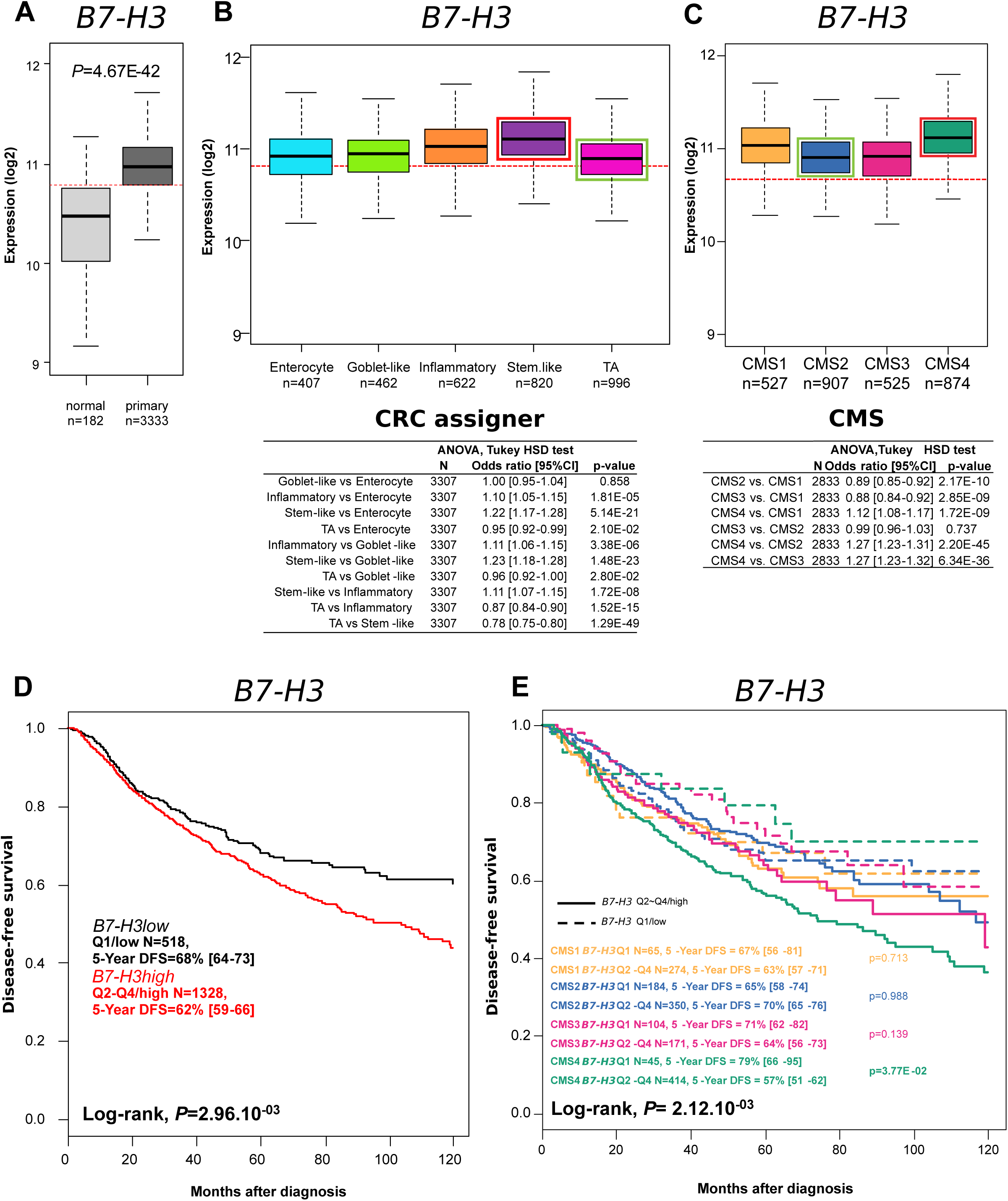

We therefore investigated the relationship between the expression level of *B7-H3* in CRC samples and clinicopathological features. CRC specimens with a high level of *B7-H3* (*B7-H3*^high^) were more frequently stage 3 (28% vs 23%, *P*=1.08 x 10^-2^), pN1 stage (32% *vs* 25%, *P*=3.09 x 10^-2^), and MSI tumors (20% vs 11%, *P*=1.62 x 10^-2^). No difference was found for patient’ age, gender, tumor location and tumor grade (Table 1). Considering the high number of cases analyzed, the clinical features between *B7-H3*^high^ and *B7-H3*^low^ CRC were overall not dramatically different.

### B7-H3 overexpression is associated with CRC molecular classifications

Several molecular classifications have been used to stratify CRC based on gene expression profiles. These classifications aim to better predict tumor outcome and therapeutic response. We therefore investigated *B7-H3* expression in the two main molecular classifications, namely the CRCassigner [32] and the Consensus Molecular Subtype (CMS) classifications [31].

The CRCassigner classification comprises 5 subclasses named according to the genes that are preferentially expressed in each of them: Goblet-like class, defined by high mRNA expression of the goblet-specific genes MUC2 and TFF3; Enterocytic class, described by high expression of enterocyte-specific genes; Stem-like class, with high expression of Wnt signaling targets as well as stem cell, myoepithelial and mesenchymal genes and low expression of differentiation markers; Inflammatory class, characterized by relative high expression of chemokines and interferon-related genes; and Transit-amplifying class, with a heterogeneous collection of samples with variable expression of stem cell and Wnt target genes. For this classification, *B7-H3^high^* tumors were enriched in the Stem-like class, while tumors with low expression of *B7-H3* were found in the Transit-amplifying class (respectively 29% and 40%, *P*=2.63 x 10^-29^) (Table 2 and Figure 1B).

The CMS classification divides CRC tumors into 4 different classes: CMS1 is characterized by a hypermutated profile, microsatellite instability (MSI) and strong immune activation; CMS2 known as the “canonical” subtype, comprises microsatellite-stable tumors with a strong epithelial phenotype and activation of Wnt and Myc signaling pathways; CMS3 is described as the “metabolic” subtype, characterized by metabolic dysregulation in epithelial cells; and finally, CMS4, the “mesenchymal” subtype, shows significant TGFβ activation, stromal invasion, and enhanced angiogenesis.

Regarding these CMS classes, *B7-H3^high^* tumors were enriched in the CMS4 class, while tumors with low expression of *B7-H3* were enriched in the CMS2 class (respectively 36% and 43%, *P*=8.43 x 10^-37^) (Table 2 and Figure 1C). Interestingly, both the Stem-like and the CMS4 classes of the two classifications are the classes with the worst prognosis at diagnosis.

### B7-H3 level of expression is associated with poor prognosis

We therefore investigated the association between *B7-H3* expression and prognosis in CRC patients, focusing on the 5-year DFS, MFS, OS. *B7-H3*^high^ was significantly associated with lower 5-year DFS and MFS (respectively 62% and 75%, *P=*2.96 x 10^-3^ and *P*=4.59 x 10^-2^), while no significant effect on 5-year OS was observed (Table 3 and Figure 1D). These results suggested that *B7-H3* expression might be a prognostic factor for relapse and progression in CRC.

In addition, we also analyzed the relationship between *B7-H3* expression and DFS within the CMS subtypes of CRC. This showed that *B7-H3*^high^ expression was significantly associated with poorer DFS, particularly in the CMS4 subtype (*P*= 2.12 x 10^-3^) (Figure 1E).

We conducted then univariate and multivariate analyzes considering clinicopathological parameters and molecular classification. In univariate analysis, patient’s age, gender, tumor location and differentiation grade showed no significant association with DFS. In contrast, tumor stage (*P*=2.95 x 10^-11^) and CMS classification (*P*=1.03 x 10^-3^) were significantly associated with DFS. Multivariate analysis integrating tumor stage and CMS classification with global *B7-H3* expression showed no significant association of *B7-H3* expression with DFS (P=0.59) (Table 4), suggesting that *B7-H3* expression is not an independent prognostic factor for DFS.

### B7-H3^high^ expression is enriched in myeloid– and fibrotic-related biological processes

We however decided to explore further the biological characteristics associated with *B7-H3* expression level in CRC, and analyzed the differentially expressed genes (DEG) between *B7-H3^high^* and the *B7-H3^low^* samples. In the training set of CRC samples (TCGA), we identified 1,798 DEG, including 1,673 genes upregulated and 125 genes downregulated in the *B7-H3^high^* CRC samples (Supplementary Table 2, Supplementary Figure S1A). This gene expression signature reliably separated samples according to *B7-H3* expression levels (*P*=5.75 × 10⁻²⁵). To validate the robustness of this signature, we tested its segregation ability according to *B7-H3* expression levels in an independent CRC cohort (COADdB), where it successfully discriminated *B7-H3^high^* and *B7-H3^low^* samples (*P*=4.49 x 10^-83^) (Supplementary Figure S1B).

The signature of *B7-H3*^high^ CRC samples was enriched with genes coding for collagens (*COL6A2, COL6A1, COL1A1, COL5A3, COL7A1, COL18A1, PCOLCE*…), laminin (*LAMB2, LAMA1*) and fibulin (*FBLN7, FBLN1, FBLN5*), fibronectin (*FN1*), which are involved in matrix assembly. It was also enriched in genes coding for matrix metalloproteinases (*MMP14, MMP23B, MMP9, MMP2*) and ADAM metallopeptidases (*ADAMTS7, ADAMTS2, ADAM8, ADAMTS10, ADAMTS4*), which are involved in matrix degradation; genes involved in cell adhesion, coding for integrins (*ITGA5, ITGB2, ITGA11, ITGB3*) were also highly expressed. Additionally, *B7-H3*^high^ CRC samples showed activation of the TGFβ pathway (*TGFB1, INHBA, TGFB3*), of the epithelial-to-mesenchymal transition (EMT) (*ZEB2, TWIST1, TWIST2*), angiogenesis (*NOTCH3, PDGFB, VEGFC, PDGFC, FGF5, ANGPTL2, ANGPTL1*), prostaglandin synthesis (*PTGS2, PTGIS, PTGDS*), hypoxia (*CA9*) and immune exhaustion (*CD274, HAVCR2, VSIR*). *B7-H3*^high^ samples also expressed an important number of CAF markers (*CAVIN1, PDGFRB, VIM, ACTA2, TAGLN, PDPN, FAP, ANTXR1, POSTN GREM1, CTHRC1*). Finally, *B7-H3*^high^ tumors were enriched in genes involved in complement cascade (*C1QC, C4A, C3, C1QA, C1QB, C7, CFD, C1R*…), antigen presentation (*HLA-DPB1, HLA-DRB1, HLA-DPA1, HLA-DRB5, CIITA*, …) and markers of the myeloid compartment, notably for type 2 macrophages (*CD14, ITGAM, ITGAX, CD163, MRC1*).

In parallel, and to strengthen these findings, we looked at the DEG ontologies (Biological Processes and Hallmarks, GSEA). We focused on the top-10 gene ontologies. *B7-H3*^high^ samples were enriched in pathways involved in adhesion (biological adhesion; *P*=2.64 x 10^-80^, cell-cell adhesion; *P*=1.41 x 10^-36^), extracellular matrix assembly (*P*=5.76 x 10^-57^), and EMT (*P*=8.65 x 10^-54^), defense response (*P*=7.54 x 10^-36^), inflammatory response (*P*=2.20 x 10^-34^) and regulation of immune system processes (*P*=1.87 x 10^-33^) (Supplementary Table 3). All these pathways were concordant with our previous analysis of DEG.

These data suggest that *B7-H3*^high^ tumors are highly fibrotic and infiltrated with immune components within an inflammatory bed.

### B7-H3*^high^* expression is associated with immune defective tumor microenvironment

To test the aforementioned enrichment in stromal and immune cells in *B7-H3*^high^ tumors, we used the Estimate score. The immune compartment was high in both group of tumors, being slightly enriched in *B7-H3*^high^ samples (*P*=2.03 × 10⁻²). The major difference between *B7-H3*^low^ and *B7-H3*^high^ samples however concerns the stromal compartment, which was more prominent in *B7-H3*^high^ tumors (*P*=6.61 × 10⁻²^0^) (Table 5 and Figure 2A).

**Figure 2.**
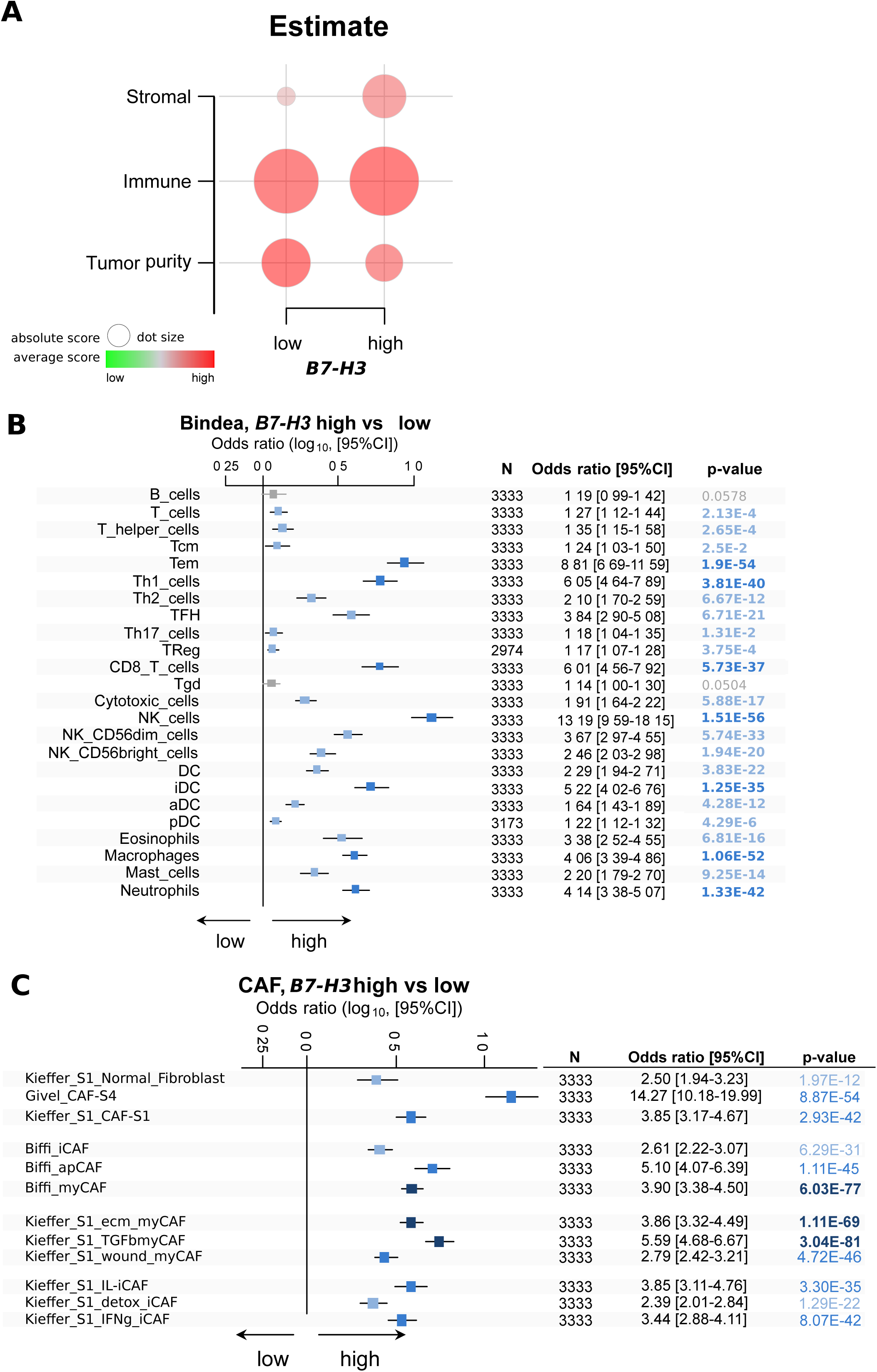

We then analyzed qualitatively the immune cells infiltrates of *B7-H3*^low^ and *B7-H3*^high^ samples by examining the association between *B7-H3* expression levels and immune cell subtypes defined by Bindea’s immunome [37]. We focused on the most significantly enriched subsets (P< 10⁻^35^), and found that *B7-H3*^high^ CRC samples were highly infiltrated with effector memory T cells (Tem; *P*=1.90 x 10^-54^), T helper 1 (Th1) cells (*P*=3.81 × 10⁻⁴⁰), CD8^+^ T cell (*P*=5.73 x 10^-37^) and NK cells (*P*=1.51 x 10^-56^). In addition, macrophages (*P*=1.06 x 10^-52^) and neutrophils (*P*=1.33 x 10^-42^) also infiltrated more strongly the *B7-H3*^high^ tumors than the B7-H3^low^ tumors (Figure 2B).

In parallel, we also tested enrichment in immune-related signatures. Notably we looked at the TLS signature [38], which is thought to indicate the presence of TILs that orchestrate an immune response. *B7-H3*^high^ CRC samples were enriched with the TLS signature (*P*=1.28 x 10^-25^). We also examined the T inflamed signature [39], which was shown to predict favorable response to immune checkpoint blockers (ICBs), and found that it was enriched in *B7-H3*^high^ CRC samples (*P*=1.73 x 10^-21^). We looked at the antigen presenting machinery score (APMS, [42] and found an enrichment in *B7-H3*^high^ CRC samples (*P*=2.55 x 10^-21^). Overall, our data suggested the presence of actors susceptible to drive an anti-tumor immune response in *B7-H3*^high^ CRC samples (Table 5).

To assess the degree of activation of this anti-tumor immune response, we then analyzed the CytoAct signature [40], which reflects the cytotoxic immune activation. *B7-H3*^high^ CRC samples showed a modest but significant enrichment in this signature (*P*=2.02 × 10⁻⁵). To further characterize the immune activation profile, we evaluated several of the Gatza’s signatures that correspond to patterns of pathway activity[41]. In line with the low activation of the CytoAct signature, the IFN-γ signature was only mildly enriched (*P*=1.64 × 10⁻²) in *B7-H3*^high^ tumors. Only the IFN-α signature was strongly enriched (*P*=7.34 × 10⁻²²). In addition, the STAT3 signaling signatures (*P*=2.25 × 10⁻²⁰), the TGFβ signature (*P*=2.08 x 10^-14^), and to a lesser extent, the hypoxia signature (*P*=1.83 x 10^-2^) were enriched in *B7-H3*^high^ tumors, suggesting pro-tumoral features. Overall, while the immune compartment in *B7-H3*^high^ CRC appears to be enriched in T cells with potential anti-tumor function, it does not display an effective anti-tumoral immune response (Table 5).

In parallel, we also wanted to specify the stromal infiltration on a qualitative level, notably regarding the fibroblast contingent, we searched for correlation between the most popular fibroblast signatures [34, 37, 45, 46] and *B7-H3* expression in CRC samples. *B7-H3*^high^ CRC showed enrichment in all subtypes of cancer-associated fibroblasts (CAFs). More precisely, *B7-H3*^high^ CRC were particularly enriched in myofibroblastic CAF (myCAF; *P*=6.03 x 10^-77^). The myCAF subset consists in CAFs specialized in extracellular matrix (ECM) assembly and remodeling, and TGFβ signaling; myCAFs are subdivided into subtypes, namely ECM-myCAF, wound-myCAF and TGFβ-myCAF that were all enriched in *B7-H3*^high^ CRC samples (*P*=1.11 x 10^-69^, *P*=4.72 x 10^-46^ and *P*=3.04 x 10^-81^, respectively) (Figure 2C).

Overall, the tumor microenvironment of *B7-H3*^high^ CRC samples appears to be infiltrated with immune cells with a cytotoxic potential against tumor cells, but whose activation is likely blunted. MyCAF cells, TGF-β signaling, and hypoxic conditions might be responsible for this. We then wondered if and how B7-H3 could be implicated in this anti-tumor response impairment by looking at which cell types express *B7-H3*.

### B7-H3 is highly expressed by cancer-associated fibroblasts in CRC

To further explore the cellular origin of *B7-H3* expression, we collected two public single-cell datasets (GSE132465, GSE144735) containing normal colon tissues and CRC samples, and examined the expression of *B7-H3* within cell clusters. We found that *B7-H3* expression was detectable in 25 to 50% of epithelial tumor cells at a weak level, and not present in normal epithelial cells (Figure 3A). *B7-H3* was however strongly expressed in 50 to 75% of cells forming the malignant stroma and not in the normal stromal compartment (Figure 3A). This expression in stromal cells was confirmed in a second dataset, namely GSE144735 (Figure 3B).

**Figure 3.**
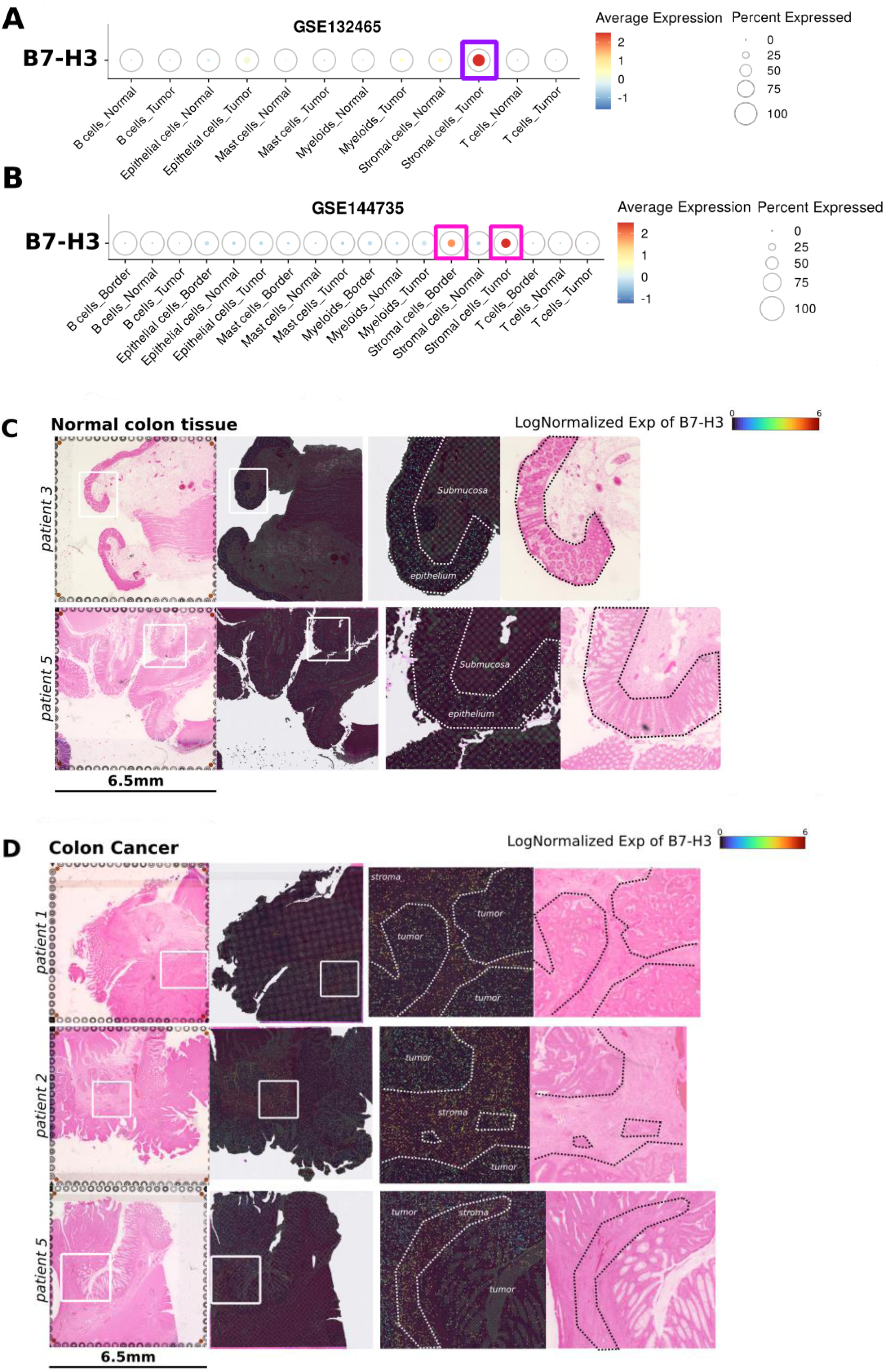

To validate these results in tissues, we analyzed spatial transcriptomic data from three primary CRC samples and two normal colon tissues (NCT). We assessed the expression of *B7-H3* and found low expression in epithelial cells of NCT. CRC tissues showed elevated *B7-H3* expression, but predominantly in the fibroblastic regions compared to epithelial tumor cells (Figure 3C-D). These results were consistent with our previous findings, but we could not exclude the possibility that post-translational modifications have altered this expression pattern. Therefore, we first investigated the correlation between *B7-H3* mRNA and protein expression in CRC cell lines. A significant correlation was observed in the CRC cell lines (r=0.65; *P*=1.56 x 10^-4^), and in all cell lines from all tumor types (r=0.76, *P*=7.21 x 10^-71^) (Supplementary Figure S1C). This suggests that *B7-H3* mRNA and B7-H3 protein expression are well correlated.

Secondly, we also validated the enrichment of *B7-H3* in fibroblasts at the protein level in tissues. To this end, we measured the expression of B7-H3, in epithelial cells (pan-KRT positive cells), and fibroblasts (pathological analysis of VIM-positive cells) in CRC tissues by multiplex immunofluorescence (N=5 patients). Interestingly, some FFPE samples contained both healthy and malignant areas. To determine the cellular expression of B7-H3, we assessed B7-H3 positivity in both compartments and confirmed that fibroblasts predominantly expressed B7-H3 compared to epithelial tumor cells. Furthermore, when comparing the healthy and tumor regions, we found that fibroblasts localized in the tumor region, *i.e.,* cancer-associated fibroblasts, overexpressed B7-H3 compared to fibroblasts in healthy region, *i.e.,* resident fibroblasts. Finally, we observed that B7-H3^high^ fibroblasts expressed less vimentin compared to adjacent B7-H3^low^ fibroblasts, indicating that only a specific subtype of fibroblasts exhibited the B7-H3^high^ phenotype (Figure 4).

**Figure 4.**
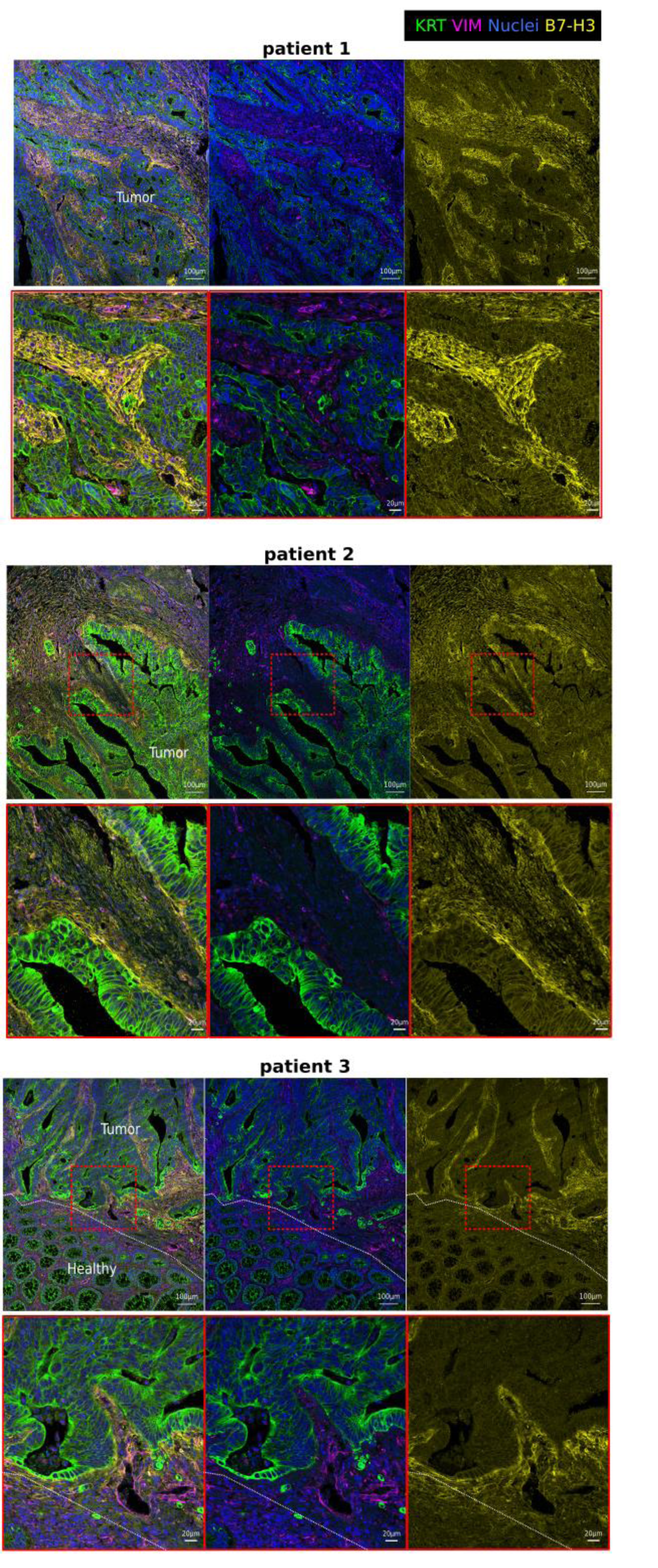

### B7-H3^high^ CAFs included two distinct subtypes

To accurately determine the nature of the *B7-H3*^high^ CAFs, we reanalyzed the two single-cell datasets previously used in Figure 3A-B. First, we re-clustered the fibroblast compartments into different subtypes (Figure 5A-B) and assessed the expression of *B7-H3* across clusters (Figure 5C-D). We then focused on clusters expressing high level of *B7-H3* (average expression >1). We then analyzed their enrichment for several reported CAF and pericyte signatures (Supplementary Table 4 and Supplementary Figure S2). In the GSE144735 dataset, cluster_c0, the cluster with the highest percentage of cells expressing a high level of *B7-H3,* showed enrichment for myCAF signatures, and cluster_c7 was enriched for the pericyte signatures (Figure 5C and E). In the GSE132465 dataset, cluster_c0, the one with the highest percentage of cells expressing a high level of *B7-H3,* was enriched in myCAF signatures; cluster_c1, was enriched in the pericyte signature; one additional cluster, cluster_c4, could be found in this dataset compared to the previous one: it was enriched in signatures related to hypoxia, Wnt signaling, and senescence pathways and was thus named *B7-H3*^high^ HWS_CAFs (Figure 5D and F). Overall, the data obtained in these two datasets are highly coherent and reproducibly identified two *B7-H3*^high^ subtypes of myCAFs and pericytes. We therefore decided to explore further the two subtypes that were common to the two datasets.

**Figure 5.**
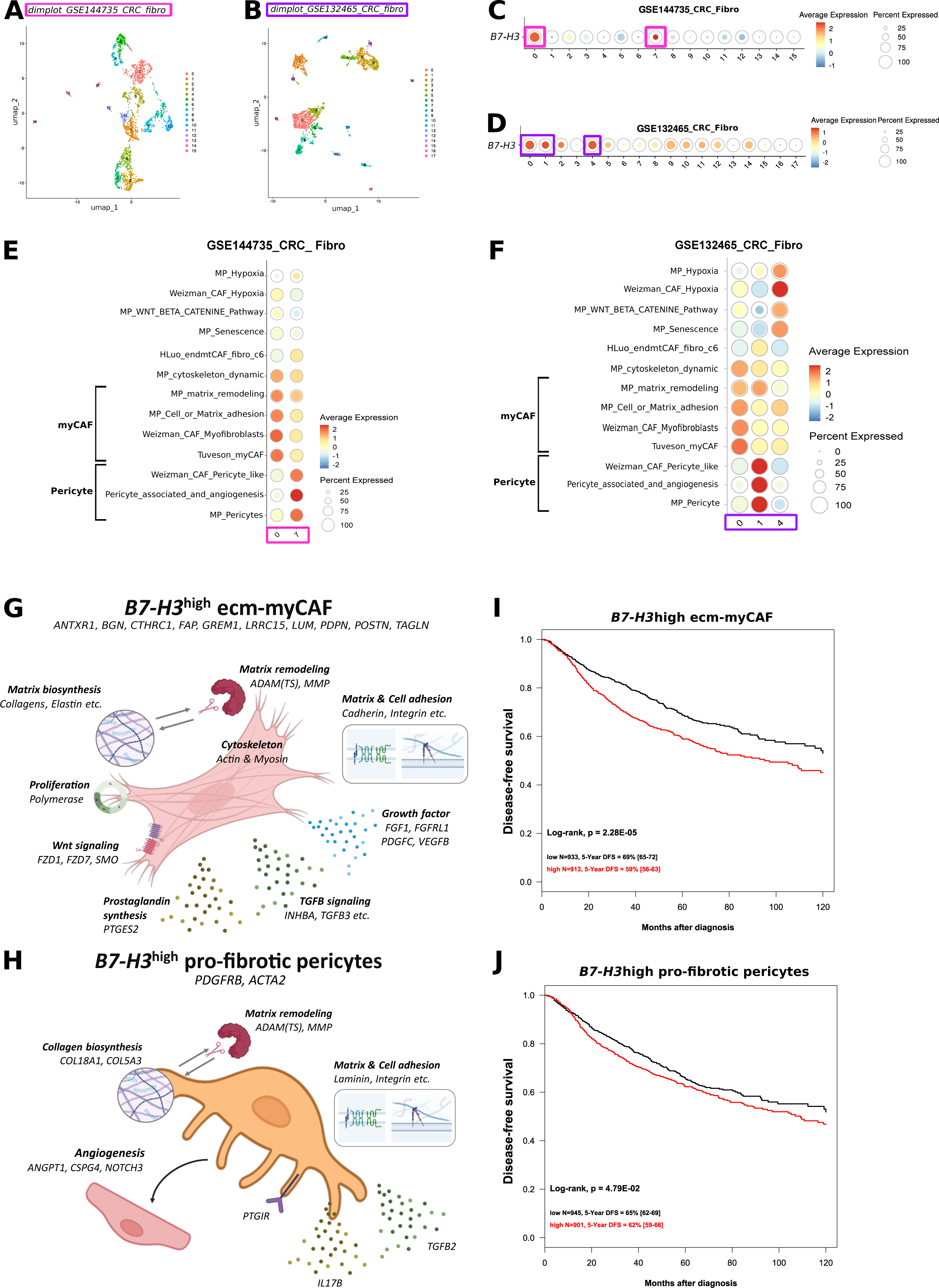

### B7-H3^high^ myCAFs are ecm_myCAF

To specify the nature of the *B7-H3*^high^ c0_myCAFs previously identified, we investigated the common characteristic genes, *i.e.*, DEGs, of the two c0_clusters and analyzed their markers and associated gene ontologies. Among the DEGs overexpressed in *B7-H3*^high^ c0_myCAF, we found genes involved in the cytoskeleton (*ACTB, ACTN1, MYH10, MYL6B, MYLK, MYO1D, MYOF*), in extracellular matrix biosynthesis (*COL10A, COL11A1, COL12A1, COL16A1, COL5A1, COL5A2, COL6A2, COL6A3, COL8A1, COL8A2, ELN, FBN1, TNC*) and remodeling *(ADAM12, ADAM19, ADAMTS12, ADAMTS2, MMP11, MMP14, MMP7, PTK7*), in CAF markers (*ANTRX1, BGN, CTHRC1, FAP, GREM1, LRRC15, LUM, PDPN, POSTN, TAGLN*), adhesion (*CDH11, CDH2, FBLN2, FN1, ICAM1, ITGA11, ITGB1, ITGB5, ITGBL1, VCAN*), growth factors (*FGF1, FGFRL1, PDGFC, VEGFB*), TGFβ pathway *(INHBA, TGFB1I1, TGFB3, TGFBI, TGFBR1*), proliferation (*POLM, POLR1E, POLR2C, POLR3D, POLR3F*), prostaglandin synthesis (*PTGES2*) (summarized in Figure 5G and detailed in Supplementary Table 5). Altogether, the DEGs indicates that *B7-H3*^high^ c0_myCAFs are strongly involved in matrix remodeling and organization. We will therefore name them *B7-H3*^high^ ecm_myCAF.

### B7-H3^high^ pericytes are pro-fibrotic pericytes

To specify the nature of *B7-H3*^high^ pericytes previously identified, we investigated the common characteristic genes, *i.e.* DEG, of the c7_cluster (GSE144735) and the c1_cluster (GSE132465) and analyzed their markers and associated gene ontologies. Among the DEGs overexpressed in *B7-H3*^high^ c7_ or c1_pericytes, we found gene involved in extracellular matrix biosynthesis (*COL18A1, COL5A3*) and remodeling (*ADAM9, ADAMTS14, ADAMTS4, MMP16*), cell and matrix adhesion (*ITGA1, ITGA4, ITGA7, LAMB2, LAMB3*), angiogenesis (*ANGPT1, CSPG4, NOTCH3*), but also markers of contractile pericytes (*ACTA2, PDGFRB*) and signaling (*IL17B, PTGIR, TGFB2*) (summarized in Figure 5H and detailed in Supplementary Table 5). Altogether, the DEGs indicates that *B7-H3*^high^ c7_ or c1_pericytes are also strongly involved in matrix remodeling and organization. We will therefore name them *B7-H3*^high^ pro-fibrosis pericytes.

### B7-H3^high^ ecm-myCAFs is an independent poor-prognosis factor in CRC patients

To investigate the involvement of these two subtypes of *B7-H3*^high^ fibroblasts (the *B7-H3*^high^ ecm_myCAFs and the *B7-H3*^high^ pro-fibrosis pericytes) on prognosis, we used the DEGs of each subtype from both datasets and selected the genes that were common in the two datasets to create two metagenes, one for each CAF subtype of interest (Supplementary Table 5). We then applied these metagenes separately to our gene expression database of bulk CRC samples to categorize patients in two groups, based on the enrichment in each metagene. We then examined the DFS in each group. Both groups enriched for each of the *B7-H3*^high^ ecm-myCAF and pro-fibrotic pericyte metagenes had lower DFS (respectively 62% *vs* 59%, *P*=4.79 x 10^-2^ and 69% vs 59%, *P=*2.28 x 10^-5^) (Figure 5I-J and Supplementary Table 6). In contrast to the multivariate analysis performed with the global expression level of *B7-H3* in bulk tissues, the multivariate analysis with *B7-H3*^high^ ecm-myCAF signature showed that *B7-H3*^high^ ecm-myCAF is an independent factor for poor prognosis in CRC patients (*P*=1.51 x 10^-2^) (Supplementary Table 5).

### B7-H3^high^ ecm_myCAF-like fibroblasts appear in the inflammatory stage of oncogenesis

We then wondered about the dynamics of appearance of *B7-H3*^High^ CAF subtypes during colon oncogenesis. To this end, we used a single-cell atlas of the colon [27] generated from cancer, normal and pre-malignant tissues, and re-clustered the fibroblast populations (Figure 6A). In order to use the nomenclature best suited to a continuum including non-malignant tissue, we will now use the term of fibroblasts rather than CAFs, which is often used in reference to malignant tissues. First, we examined the expression of *B7-H3* in fibroblast subtypes at each stage of oncogenesis. We focused in clusters expressing predominantly and strongly B7-H3 and found three subtypes of fibroblasts: cluster_c5, _c10 and _c13. The expression of *B7-H3* in cluster_c13 was also present in inflamed tissues, but not in cluster_c5 and cluster_c10, suggesting a distinct pattern of *B7-H3* expression in these subtypes (Figure 6B).

**Figure 6.**
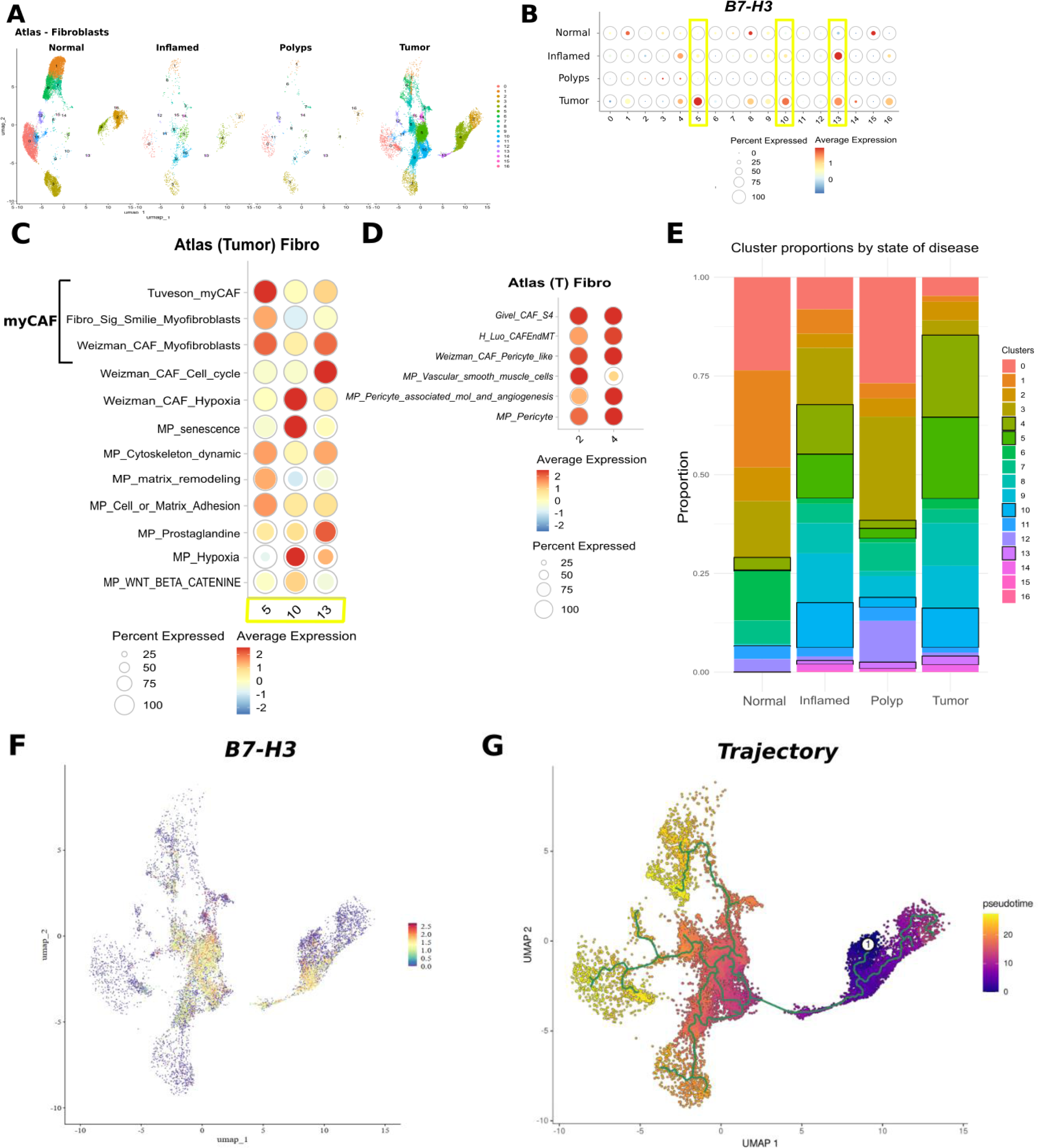

We focused on these 3 clusters and wanted to characterize them. Similar to Figure 5, we tested pathways enrichment for several fibroblast subtypes and pericyte signatures. We found that cluster_c5 activated the same pathways as the ecm-myCAFs subtype previously described; cluster_c10 activated the same pathways (hypoxia, senescence) as the HWS_CAFs cluster_c4 described in Figure 5F; cluster_c13 was however slightly distinct from the previously described fibroblast subsets, showing a proliferative profile and activation of the prostaglandin pathway (Figure 6C).

With regards to results obtained in Figure 5, we also looked for pericytes expressing *B7-H3*. We found two clusters of pericytes, cluster c2 and c4. Cluster_c2 and cluster_c4 were both enriched in pericytes signatures but cluster_c2 was also enriched in vascular smooth muscle cells (VSMC), while cluster_c4 was strongly enriched in an endo-mesenchymal transition (EndMT) signature (Figure 6D). Cluster_c4 expressed *B7-H3* at an average level in inflamed and cancerous tissues, whereas cluster_c2 did not express *B7-H3* at all (Figure 6B).

Overall, these data suggested that fibroblast subsets with similar DEG and function to the *B7-H3*^high^ ecm-myCAF, HWS_CAFs and *B7-H3*^high^ pro-fibrosis pericytes are present in non-malignant samples, but they do not express high level of *B7-H3* at these stages.

Furthermore, beside cancerous tissues, clusters with high percentage of fibroblasts expressing *B7-H3* were also found in inflamed tissues (clusters_c4, c13 and to a lesser level c5 and c10), but not in the normal tissues or polyps, suggesting that inflammation might play a role in its regulation.

To test this hypothesis, we examined the presence of these clusters of fibroblasts and look at their respective proportion during the continuum to CRC progression. Cluster_c13 showed progressive accumulation during oncogenesis and peaked at the tumor stage (FDR=8.75 × 10⁻⁹). Cluster_c5 and cluster_c4 were absent in normal colon and accumulated in inflamed colon tissue and even more in tumor samples (respectively, FDR=8.38 × 10⁻⁹ and FDR=732.20 x 10^-24^) but was only slightly present in polyps. Similarly, cluster_c10 was absent in normal colon but increased in both inflamed and tumor samples (FDR=413.86 × 10⁻¹^2^) (Figure 6E and Supplementary Table 7). These results suggested that *B7-H3*^high^ subsets (clusters_c4, _c5, _c10, and _c13) with different functions can be detected as early as the inflammatory pre-cancerous stage, and are potentially triggered by inflammation.

To establish if a differentiation continuum might exist between these subsets of fibroblasts exist, we inferred the trajectory of fibroblasts clusters using pseudotime analysis with cluster_c4 as origin. We found a consistent trajectory from cluster_c2 *B7-H3*^negativ^ to _c4 *B7-H3*^mid^ pericytes, then to cluster_c13 B7-H3^high^ proliferative fibroblasts that reach to cluster _c10 B7-H3^high^ HWS_CAF then to cluster_c5 B7-H3^superhigh^ ecm-myCAF (Figure 6F-G). Furthermore, cluster_c4 exhibited a heterogeneous expression of B7-H3 that was consistent with previous Figure 6C. Altogether, these data suggest the existence of a polarization continuum from B7-H3^neg^ pericytes to B7-H3^high^ ecm-myCAF.

## Discussion

In this study, we confirmed that B7-H3 is enriched in CRC compared to normal colon tissue, especially in tumors with poor prognosis for DFS. Samples with high levels of *B7-H3* were more fibrotic and enriched with anti-tumor immune cells that are poorly activated. More importantly, the strongest expression of *B7-H3* was observed in the fibroblast compartment, rather than in the epithelial cells in CRC samples. Several subtests of fibroblasts expressed *B7-H3*, and at least two were redundant in several datasets highlighting their robustness: the pro-fibrotic pericytes and the ecm-myCAF. We showed that the *B7-H3*^high^ ecm-myCAF subset was an independent poor-prognosis for DFS, that shared some similarities with fibroblasts detected in the pre-cancerous inflamed stage during oncogenesis. Interestingly, CAF-mediated ECM remodeling and fibrosis have been reported to contribute to the formation of an immunosuppressed microenvironment that promotes tumor growth [34, 47]. In addition to a physical barrier, we have found that our ecm-myCAFs produced members of the PGE2 and TGFβ families that are known to play an immunosuppressive role on the anti-tumor immunity, and simultaneously encourage tumor cells proliferation. Our study revealed that *B7-H3^high^* ecm-myCAF may be a promising therapeutic target with a double benefit (improved anti-tumor response and decrease tumor cell proliferation) for CRC patients.

The immune function of B7-H3 has long been controversial, fluctuating between a role as a co-stimulatory or immunosuppressive molecule [48–51] and its receptor is still unknown. One explanation for these apparently contradictory functions could be that B7-H3, or its receptor(s), have multiple expression patterns, allowing it to fulfill multiple functions. Here, we have shown that B7-H3 were enriched in two classes of CRC; the immune-related CMS1 and the mesenchymal-related CMS4, both presenting diametrically distinct malignant specificities. We also showed that *B7-H3*^high^ samples were enriched in myeloid cells. Functional and deeper study of the interaction of *B7-H3*^high^ ecm_myCAF with other cells from the tumor ecosystem could allow to precise the mechanistic underlined, for example the setting of a myeloid-based immunosuppressive microenvironment. But this will require a better knowledge of the receptor(s) for B7-H3.

Now that our results shed light on the *B7-H3*^high^ ecm-CAF subsets with prognostic features, a deeper functional investigation might be warranted. In this line, CAFs expressing B7-H3 protein were also found in renal cell carcinomas. Silencing B7-H3 in these CAFs reduced their proliferation and induced their apoptosis [52]. In fibroblasts cocultured with renal cancer cells, B7-H3 knock-down decreased the proliferation and migratory capacity of tumor cells, which was due to reduced secretion of HGF and CXCL12. Overall, silencing of B7-H3 in fibroblasts led to reduced tumor size in xenograft [52]. The myCAF polarization of these fibroblasts was not further explored in this study. Nevertheless, this study confirmed the presence of B7-H3 expressing CAFs in other cancers and confirmed their pro-tumoral behavior. Extending our approach to other solid tumor datasets may allow us to reinforce the interest in targeting B7-H3 ecm-myCAF and investigate tissue-specific subtleties.

Interestingly, we reported on the chronology of *B7-H3*^high^ fibroblast subsets occurrence during oncogenesis. However, this needs to be confirmed in other datasets, when available, as indeed, for now datasets gathering tissue of interest and of sufficient quality remain scarce. Notably in our case, the number of cells from the polyp stage is particularly low. Similarly, we also analyzed metastatic tissues of colon cancer (Supplementary Figure S3A-C). We found only one cluster expressing high level of B7-H3, which was enriched for myCAF markers, but also in inflammation signature and WNT pathway, suggesting potential changes in metastatic stage compared to primary CRC. Here again, the quality of the data limited the interpretation. In a near future, single nuclei-data, which are more appropriate for the analysis of the fibroblast compartment than single cell data, of pre-cancerous and advanced malignant stages will be available to confirm our findings.

Nevertheless, with the available datasets, we found that a subset of fibroblasts expressing high levels of *B7-H3* could be detected in inflamed colon tissue, maintained in tumor tissue, but was absent in normal colon. This suggests that inflammation could promote the differentiation and/or expansion of *B7-H3*^high^ fibroblast subsets, lying the pre-malignant bed. This possibility remains speculative and deserves further investigation about their role as a potential predictive marker for CRC development.

In the meantime, several trials are investigating the impact of B7-H3 targeting in oncology. Currently, the most advanced anti-B7-H3 therapy is Ifinatamab Deruxtecan (I-DXd, also known as DS-7300), an antibody-drug conjugate to a DNA topoisomerase I, developed by Merck and Daiichi Sankyo™. Several clinical trials from phase IB to phase III are ongoing in advanced and metastatic solid tumors (NCT06330064, NCT04145622), notably small cell lung cancer (NCT05280470, NCT06362252, NCT06203210, NCT06780137), prostate cancer (NCT06863272, NCT06925737) and esophageal squamous cell carcinoma (NCT06644781a) to investigate the benefit of targeting B7-H3 in all tumor tissues.

Overall, our results confirm the tumor-promoting role of B7-H3 in CRC and justify its targeting. However, we clarify its expression in CAFs, and especially in ecm-myCAFs with poor prognosis value for DFS. Based on others and our reports, using an ADC to target B7-H3 implies that cells from the microenvironment, and especially myCAFs, will be primarily eliminated. This introduces a new concept of targeting the supporting niche rather than the tumor cells directly. The results of the current clinical trials based on an anti-B7-H3 ADC will therefore be particularly interesting to interpret with these data in mind.

## Author Contributions

Conception and design, E.M., D.J.B., F.B.; methodology, E.M., M.P., BdR; validation, E.M.; formal analysis, P.F., M.P., A.G., N.B., B.D.R., L.M., E.M.; resources, L.M., N.B., B.D.R, D.J.B; data curation, M.P., A.G., P.F., L.M.; writing—original draft preparation, M.P., E.M., A.G., P.F., N.B., B.D.R.; writing—review and editing, M.P., E.M., F.B., P.F, AG., supervision, E.M. All authors have read and agreed to the published version of the manuscript.

## Funding

Ligue contre le Cancer – Bertucci

Cancéropole PACA

GIRCI Mediterranée

## Institutional Review Board Statement

The study was approved by our institutional review board.

## Informed Consent Statement

Our in-silico study is based on public data from published studies in which the informed patient’s consent to participate and the ethics and institutional review board were already obtained by authors. Our in vitro study is based on a prospective cohort for which informed consent availability is an inclusion criteria.

## Data Availability Statement

This study did not generate any new material. All links or accession numbers are listed in the Materials and Methods section.

## Conflicts of Interest

The authors declare no conflict of interest.

## Acknowledgment

The authors wish to thank Dr. Caroline Gouarne and Dr. Brice Chanez for their contribution in the CTC-Colon cohort that provide FFPE samples and the IPC/CRCM Experimental Pathology Plateform (ICEP) and MICS plateform for their technical support.

## Notes

### Competing Interest Statement

The authors have declared no competing interest.

## Bibliography

1. Sung H, Ferlay J, Siegel RL, et al. Global Cancer Statistics 2020: GLOBOCAN Estimates of Incidence and Mortality Worldwide for 36 Cancers in 185 Countries. CA Cancer J Clin 2021;71(3):209–249.

2. Lecomte T, Tougeron D, Chautard R, et al. Non-metastatic colon cancer: French Intergroup Clinical Practice Guidelines for diagnosis, treatments, and follow-up (TNCD, SNFGE, FFCD, GERCOR, UNICANCER, SFCD, SFED, SFRO, ACHBT, SFP, AFEF, and SFR). Dig Liver Dis 2024;56(5):756–769.

3. Siegel RL, Wagle NS, Cercek A, et al. Colorectal cancer statistics, 2023. CA Cancer J Clin 2023;73(3):233–254.

4. Dekker E, Tanis PJ, Vleugels JLA, et al. Colorectal cancer. Lancet 2019;394(10207):1467–1480.

5. Jaime-Casas S, Barragan-Carrillo R, Tripathi A. Antibody-drug conjugates in solid tumors: a new frontier. Curr Opin Oncol 2024;36(5):421–429.

6. Tarantino P, Carmagnani Pestana R, Corti C, et al. Antibody-drug conjugates: Smart chemotherapy delivery across tumor histologies. CA Cancer J Clin 2022;72(2):165–182.

7. Roth TJ, Sheinin Y, Lohse CM, et al. B7-H3 ligand expression by prostate cancer: a novel marker of prognosis and potential target for therapy. Cancer Res 2007;67(16):7893–900.

8. Brunner A, Hinterholzer S, Riss P, et al. Immunoexpression of B7-H3 in endometrial cancer: relation to tumor T-cell infiltration and prognosis. Gynecol Oncol 2012;124(1):105–11.

9. Flem-Karlsen K, Tekle C, Andersson Y, et al. Immunoregulatory protein B7-H3 promotes growth and decreases sensitivity to therapy in metastatic melanoma cells. Pigment Cell Melanoma Res 2017;30(5):467–476.

10. Zang X, Sullivan PS, Soslow RA, et al. Tumor associated endothelial expression of B7-H3 predicts survival in ovarian carcinomas. Mod Pathol 2010;23(8):1104–12.

11. Sun Y, Wang Y, Zhao J, et al. B7-H3 and B7-H4 expression in non-small-cell lung cancer. Lung Cancer 2006;53(2):143–51.

12. Zhao X, Li DC, Zhu XG, et al. B7-H3 overexpression in pancreatic cancer promotes tumor progression. Int J Mol Med 2013;31(2):283–91.

13. Wang R, Ma Y, Zhan S, et al. B7-H3 promotes colorectal cancer angiogenesis through activating the NF-κB pathway to induce VEGFA expression. Cell Death Dis 2020;11(1):55.

14. Bin Z, Guangbo Z, Yan G, et al. Overexpression of B7-H3 in CD133+ colorectal cancer cells is associated with cancer progression and survival in human patients. J Surg Res 2014;188(2):396–403.

15. Jiang B, Zhang T, Liu F, et al. The co-stimulatory molecule B7-H3 promotes the epithelial-mesenchymal transition in colorectal cancer. Oncotarget 2016;7(22):31755–71.

16. Liu F, Zhang T, Zou S, et al. B7-H3 promotes cell migration and invasion through the Jak2/Stat3/MMP9 signaling pathway in colorectal cancer. Mol Med Rep 2015;12(4):5455–60.

17. Mao Y, Chen L, Wang F, et al. Cancer cell-expressed B7-H3 regulates the differentiation of tumor-associated macrophages in human colorectal carcinoma. Oncol Lett 2017;14(5):6177–6183.

18. Lu Z, Zhao ZX, Cheng P, et al. B7-H3 immune checkpoint expression is a poor prognostic factor in colorectal carcinoma. Mod Pathol 2020;33(11):2330–2340.

19. Ingebrigtsen VA, Boye K, Tekle C, et al. B7-H3 expression in colorectal cancer: nuclear localization strongly predicts poor outcome in colon cancer. Int J Cancer 2012;131(11):2528–36.

20. Wang W, Li Q, Yamada T, et al. Crosstalk to stromal fibroblasts induces resistance of lung cancer to epidermal growth factor receptor tyrosine kinase inhibitors. Clin Cancer Res 2009;15(21):6630–8.

21. Gaggioli C, Hooper S, Hidalgo-Carcedo C, et al. Fibroblast-led collective invasion of carcinoma cells with differing roles for RhoGTPases in leading and following cells. Nat Cell Biol 2007;9(12):1392–400.

22. Biffi G, Tuveson DA. Diversity and Biology of Cancer-Associated Fibroblasts. Physiol Rev 2021;101(1):147–176.

23. Cui JY, Ma J, Gao XX, et al. Unraveling the role of cancer-associated fibroblasts in colorectal cancer. World J Gastrointest Oncol 2024;16(12):4565–4578.

24. Özdemir BC, Pentcheva-Hoang T, Carstens JL, et al. Depletion of carcinoma-associated fibroblasts and fibrosis induces immunosuppression and accelerates pancreas cancer with reduced survival. Cancer Cell 2014;25(6):719–34.

25. Lee HO, Hong Y, Etlioglu HE, et al. Lineage-dependent gene expression programs influence the immune landscape of colorectal cancer. Nat Genet 2020;52(6):594–603.

26. Che LH, Liu JW, Huo JP, et al. A single-cell atlas of liver metastases of colorectal cancer reveals reprogramming of the tumor microenvironment in response to preoperative chemotherapy. Cell Discov 2021;7(1):80.

27. Chu X, Li X, Zhang Y, et al. Integrative single-cell analysis of human colorectal cancer reveals patient stratification with distinct immune evasion mechanisms. Nat Cancer 2024;5(9):1409–1426.

28. Taminau J, Steenhoff D, Coletta A, et al. inSilicoDb: an R/Bioconductor package for accessing human Affymetrix expert-curated datasets from GEO. Bioinformatics 2011;27(22):3204–5.

29. Bertucci F, Finetti P, Viens P, et al. EndoPredict predicts for the response to neoadjuvant chemotherapy in ER-positive, HER2-negative breast cancer. Cancer Lett 2014;355(1):70–5.

30. Johnson WE, Li C, Rabinovic A. Adjusting batch effects in microarray expression data using empirical Bayes methods. Biostatistics 2007;8(1):118–27.

31. Guinney J, Dienstmann R, Wang X, et al. The consensus molecular subtypes of colorectal cancer. Nat Med 2015;21(11):1350–6.

32. Sadanandam A, Lyssiotis CA, Homicsko K, et al. A colorectal cancer classification system that associates cellular phenotype and responses to therapy. Nat Med 2013;19(5):619–25.

33. Öhlund D, Handly-Santana A, Biffi G, et al. Distinct populations of inflammatory fibroblasts and myofibroblasts in pancreatic cancer. J Exp Med 2017;214(3):579–596.

34. Kieffer Y, Hocine HR, Gentric G, et al. Single-Cell Analysis Reveals Fibroblast Clusters Linked to Immunotherapy Resistance in Cancer. Cancer Discov 2020;10(9):1330–1351.

35. Li H, Courtois ET, Sengupta D, et al. Reference component analysis of single-cell transcriptomes elucidates cellular heterogeneity in human colorectal tumors. Nat Genet 2017;49(5):708–718.

36. Yoshihara K, Shahmoradgoli M, Martínez E, et al. Inferring tumour purity and stromal and immune cell admixture from expression data. Nat Commun 2013;4:2612.

37. Bindea G, Mlecnik B, Tosolini M, et al. Spatiotemporal dynamics of intratumoral immune cells reveal the immune landscape in human cancer. Immunity 2013;39(4):782–95.

38. Coppola D, Nebozhyn M, Khalil F, et al. Unique ectopic lymph node-like structures present in human primary colorectal carcinoma are identified by immune gene array profiling. Am J Pathol 2011;179(1):37–45.

39. Ayers M, Lunceford J, Nebozhyn M, et al. IFN-γ-related mRNA profile predicts clinical response to PD-1 blockade. J Clin Invest 2017;127(8):2930–2940.

40. Rooney MS, Shukla SA, Wu CJ, et al. Molecular and genetic properties of tumors associated with local immune cytolytic activity. Cell 2015;160(1-2):48–61.

41. Gatza ML, Lucas JE, Barry WT, et al. A pathway-based classification of human breast cancer. Proc Natl Acad Sci U S A 2010;107(15):6994–9.

42. Thompson JC, Davis C, Deshpande C, et al. Gene signature of antigen processing and presentation machinery predicts response to checkpoint blockade in non-small cell lung cancer (NSCLC) and melanoma. J Immunother Cancer 2020;8(2).

43. Hochberg Y, Benjamini Y. More powerful procedures for multiple significance testing. Stat Med 1990;9(7):811–8.

44. Andreatta M, Carmona SJ. UCell: Robust and scalable single-cell gene signature scoring. Comput Struct Biotechnol J 2021;19:3796–3798.

45. Givel AM, Kieffer Y, Scholer-Dahirel A, et al. miR200-regulated CXCL12β promotes fibroblast heterogeneity and immunosuppression in ovarian cancers. Nat Commun 2018;9(1):1056.

46. Elyada E, Bolisetty M, Laise P, et al. Cross-Species Single-Cell Analysis of Pancreatic Ductal Adenocarcinoma Reveals Antigen-Presenting Cancer-Associated Fibroblasts. Cancer Discov 2019;9(8):1102–1123.

47. Croizer H, Mhaidly R, Kieffer Y, et al. Deciphering the spatial landscape and plasticity of immunosuppressive fibroblasts in breast cancer. Nat Commun 2024;15(1):2806.

48. Koumprentziotis IA, Theocharopoulos C, Foteinou D, et al. New Emerging Targets in Cancer Immunotherapy: The Role of B7-H3. Vaccines (Basel) 2024;12(1).

49. Yi KH, Chen L. Fine tuning the immune response through B7-H3 and B7-H4. Immunol Rev 2009;229(1):145–51.

50. Chapoval AI, Ni J, Lau JS, et al. B7-H3: a costimulatory molecule for T cell activation and IFN-gamma production. Nat Immunol 2001;2(3):269–74.

51. Loos M, Hedderich DM, Friess H, et al. B7-h3 and its role in antitumor immunity. Clin Dev Immunol 2010;2010:683875.

52. Zhang S, Zhou C, Zhang D, et al. The anti-apoptotic effect on cancer-associated fibroblasts of B7-H3 molecule enhancing the cell invasion and metastasis in renal cancer. Onco Targets Ther 2019;12:4119–4127.

