## Supplementary material for "Multi-omic analysis of colorectal adenocarcinoma identifies a new subtype of myofibroblastic cancer-associated fibroblast expressing high level of B7-H3 and with poor prognosis value": Legends

**Figure 1**: Expression and prognostic factor of *B7-H3* expression in CRC samples

(A) Box plot representing *B7-H3* expression in normal colon tissue compared to primary CRC tumor in bulk primary CRC dataset. Significance between groups was defined by Student t-test.
(B) Box plot representing *B7-H3* expression in CRCassigner subclassification of CRC samples in bulk primary CRC dataset. Red frame highlights the CRCassigner subclass the most enriched in *B7-H3* expression whereas the green one highlights the CRCassigner subclass the less enriched in *B7-H3* expression. Significance was defined using an ANOVA Tukey HSD test.
(C) Box plot representing *B7-H3* expression in CMS subclassification of CRC samples in bulk primary CRC dataset. Red frame highlights the CMS subclass the most enriched in *B7-H3* expression whereas the green one highlights the CMS subclass the less enriched in *B7-H3* expression. Significance was obtained using an ANOVA Tukey HSD test.
(D) Kaplan-Meier survival curve representing the disease-free survival of B7-H3$^{high}$ (red) group and B7-H3$^{low}$ (black) group in regards to time (months after diagnosis). Significance was defined using a log-rank test.
(E) Kaplan-Meier survival curve representing the disease-free survival according to CMS subclasses (CMS1 in yellow, CMS2 in blue, CMS3 in pink and CMS4 in green) segregated between B7-H3$^{high}$ (solid line) and B7-H3$^{low}$ (dash line), in regards to time (months after diagnosis). Significance was defined using a log-rank test.

**Figure 2:** Enrichment of tumor microenvironment features in regards to B7-H3 level of expression

(A) Bubble plot representing average score of Estimate classification according B7-H3$^{high}$ and B7-H3$^{low}$ groups in bulk dataset. Enrichment was represented in relative value of average score level using green-to-red scale and in absolute value using the size of the circle. Score range and P-value were obtained using Student t-test.
(B) Forest plot representing the enrichment of immunome (Bindea *et al.*) between B7-H3$^{high}$ and B7-H3$^{low}$ groups. P-value and odd-ratio were represented in adjacent table. Significant enrichments were represented in blue, darkest being the most significant.
(C) Forest plot representing the enrichment of cancer-associated signatures (CAF) between B7-H3$^{high}$ and B7-H3$^{low}$ groups. P-value and odd-ratio were represented in adjacent table. Significant enrichments were represented in blue, darkest being the most significant.

**Figure 3:** B7-H3 expression in epithelial and stromal cells in CRC and normal adjacent tissue

(A) Dot plot representing the expression of *B7-H3* in cell subtypes of a single-cell dataset (GSE132465) of normal tissue and CRC samples. Circle size represented the percentage of cell enriched in B7-H3 and the colored scale represented the average expression. Most enriched clusters were framed.
(B) Dot plot representing the expression of B7-H3 in cell subtypes of a single-cell dataset (GSE144735) of normal tissue and CRC samples. Circle size represented the percentage of cell enriched in B7-H3 and the colored scale represented the average expression. Most enriched clusters were framed.
(C) Histologic and projection of B7-H3 expression of *B7-H3* in spatial transcriptomic data of Normal tissue (N=2 patients). Colored-scale represents the level of expression. Epithelial and stroma region were defined in dash line.

(D) Histologic and projection of B7-H3 expression in spatial transcriptomic data of CRC (N=3 patients). Colored-scale represents the level of expression. Tumor and stroma region were defined in das line.

**Figure 4:** Expression of B7-H3 at the protein level in CRC tissues. Immunofluorescence pictures of B7-H3 expression in CRC tissues. The whole tissue and an insert were shown. B7-H3 was represented in yellow, fibroblast in pink (VIM), tumor in green (KRT) and nuclei in blue (DAPI). Healthy and tumoral region were defined when needed by dash line.

**Figure 5:** Nature and prognostic value of B7-H3$^{high}$ fibroblasts in CRC samples

(A) uMAP dimplot representation of fibroblast clusters sub-clusterisation of GSE144735 scRNAseq dataset.
(B) uMAP dimplot representation of fibroblast clusters sub-clusterisation of GSE132465 scRNAseq dataset.
(C) Dotplot representation of *B7-H3* expression in GSE144735 fibroblasts subclusters. Circle size represented the percentage of cell enriched in *B7-H3* and the color represented the average expression. Most enriched clusters were framed.
(D) Dotplot representation of *B7-H3* expression in GSE312465 fibroblasts subclusters. Circle size represented the percentage of cell enriched in *B7-H3* and the color represented the average expression. Most enriched clusters were framed.
(E) Dotplot representation of several CAF-related signatures expression in GSE144735 B7-H3$^{high}$ fibroblasts subclusters. Circle size represented the percentage of cell enriched in each signature and the color represented the average expression.
(F) Dotplot representation of several CAF-related signatures expression in GSE132465 B7-H3$^{high}$ fibroblasts subclusters. Circle size represented the percentage of cell enriched in each signature and the color represented the average expression.
(G) Summary illustration of B7-H3$^{high}$ ecm-myCAF main characteristics. Made with Biorender.
(H) Summary illustration of B7-H3$^{high}$ pro-fibrotic pericytes main characteristics. Made with Biorender.
(I) Kaplan-Meier survival curve representing the disease-free survival between CRC samples enriched in the B7-H3$^{high}$ ecm-myCAF metagene (red), in regards to time (months after diagnosis), or not (black). Significance was defined using Log-rank.
(J) Kaplan-Meier survival curve representing the disease-free survival between CRC samples enriched in the B7-H3$^{high}$ pro-fibrotique pericyte metagene (red) in regards to time (months after diagnosis), or not (black). Significance was defined using Log-rank. CRC samples enriched in the metagene was represented in red.

**Figure 6:** Nature of B7-H3$^{high}$ fibroblast across CRC oncogenesis

(A) Serial uMAP dimplots representing fibroblast sub-clusterisations in the atlas dataset (GSE161277) depending on oncogenesis status (Normal, Inflamed, Polyps and Tumor).
(B) Dotplot representing *B7-H3* expression of fibroblasts subclusters depending on oncogenesis status. Circle size represented the percentage of cell enriched in each signature and the colored scale represented the average expression. Most enriched clusters were framed.
(C) Dotplot representation of several CAF-related signatures expression in the Atlas B7-H3$^{high}$ fibroblast subclusters. Circle size represented the percentage of cell enriched in each signature and the colored scale represented the average expression.

(D) Dotplot representation of pericytes-related signatures expression in the Atlas B7-H3^high fibroblast subclusters. Circle size represented the percentage of cell enriched in each signature and the colored scale represented the average expression.
(E) Proportion of fibroblast subclusters of Atlas dataset over oncogenesis status (Normal, Inflamed, Polyps and Tumor). *B7-H3*^high fibroblasts were framed.
(F) Dimplot representation of *B7-H3* expression in fibroblasts clusters of CRC samples. The colored scale represented the average expression.
(G) Trajectory inference of fibroblasts clusters, with cluster_c4 defined as origin. The colored-scale represented the proximity between clusters.

**Supplementary Figure S1**: Supervised analysis of gene expression profiles between the "*B7-H3*-high" and "*B7-H3*-low" classes and correlation of *B7-H3* protein with RNA expression

(A) Volcano plot representing the 1,798 genes differentially expressed in the learning set (TCGA) between B7-H3^high and ^low groups. Red genes are up-regulated in B7-H3^high samples whereas green genes are up-regulated in B7-H3^low samples. Significant genes were defined by a moderated t-test p<5% q<5% and fold change > 1.5x.
(B) Box plot representing metagene prediction scores of *B7-H3* status in the learning set (left) and in validation (right).
(C) Correlation between protein and transcript expression of *B7-H3* in all CCLE cell lines (grey) and in colon cancer cell lines (blue). CL means cell line.

**Supplementary Figure S2:** All CAF signatures tested on B7-H3^high fibroblasts

(A) Dotplot representation of the expression of all CAF-related signatures tested in the GSE144735 B7-H3^high fibroblasts subclusters. Circle size represented the percentage of cell enriched in each signature and the colored scale represented the average expression.
(B) Dotplot representation of the expression of all CAF-related signatures tested in the GSE132465 B7-H3^high fibroblasts subclusters. Circle size represented the percentage of cell enriched in each signature and the colored scale represented the average expression.
(C) Dotplot representation of the expression of all CAF-related signatures tested in the Atlas B7-H3^high fibroblasts subclusters. Circle size represented the percentage of cell enriched in each signature and the colored scale represented the average expression.

**Supplementary Figure S3:** Nature of B7-H3^high fibroblast in metastatic CRC

(A) uMAP dimplot representation of fibroblast subclusters in a metastatic CRC scRNAseq dataset (GSE178318).
(B) Dot plot representing *B7-H3* expression of metastatic fibroblasts subclusters. Circle size represented the percentage of cell enriched in each signature and the colored scale represented the average expression. The most enriched cluster was framed.
(C) Dotplot representation of several CAF-related signatures expression in the metastatic CRC B7-H3^high fibroblast subcluster. Circle size represented the percentage of cell enriched in each signature and the colored scale represented the average expression.
