## Supplemental Figure 1 for "Multi-omic analysis of colorectal adenocarcinoma identifies a new subtype of myofibroblastic cancer-associated fibroblast expressing high level of B7-H3 and with poor prognosis value"

### Supplementary Figure 1

**A** **Supervised analysis**
TCGA, Learning set

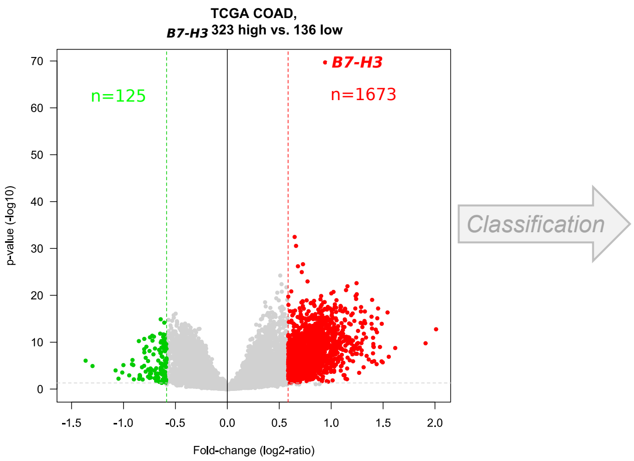


**B**

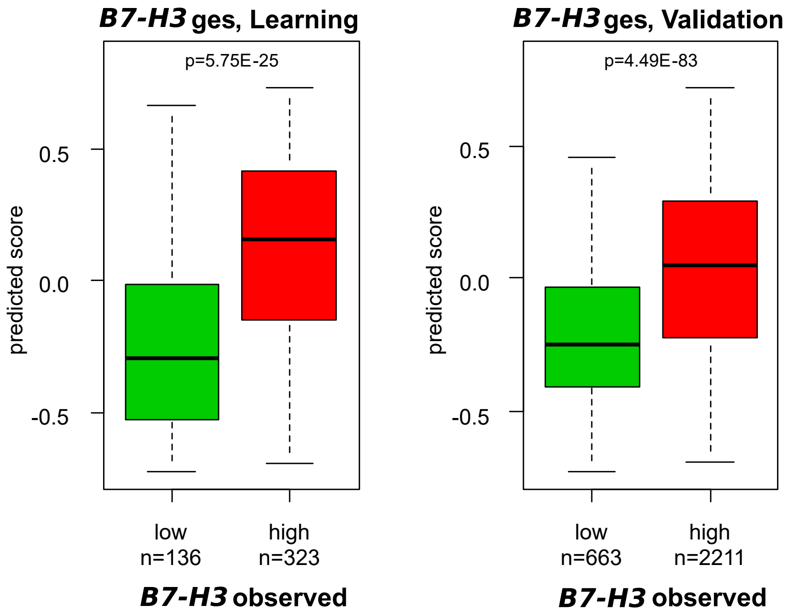


**C**

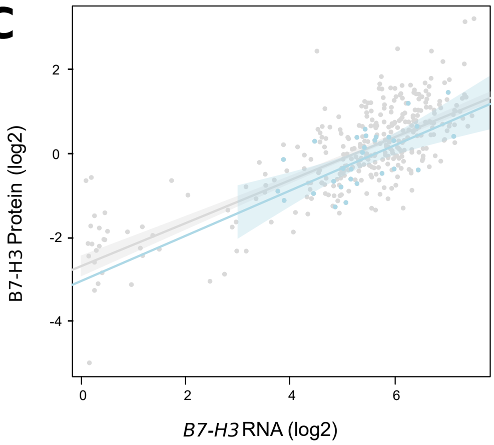


| DepMap, RNA ~ Prot, all CL | N | Pearson, r | p-value |
|---|---|---|---|
| *B7-H3* | 663 | 0.76 | 7.21E-71 |

| DepMap, RNA ~ Prot, Colorectal CL | N | Pearson, r | p-value |
|---|---|---|---|
| *B7-H3* | 29 | 0.65 | 1.56E-04 |
