## Supplementary figures and images for "Multi-omic analysis of colorectal adenocarcinoma identifies a new subtype of myofibroblastic cancer-associated fibroblast expressing high level of B7-H3 and with poor prognosis value"

### Supplemental Figure 2

#### Supplementary Figure 2

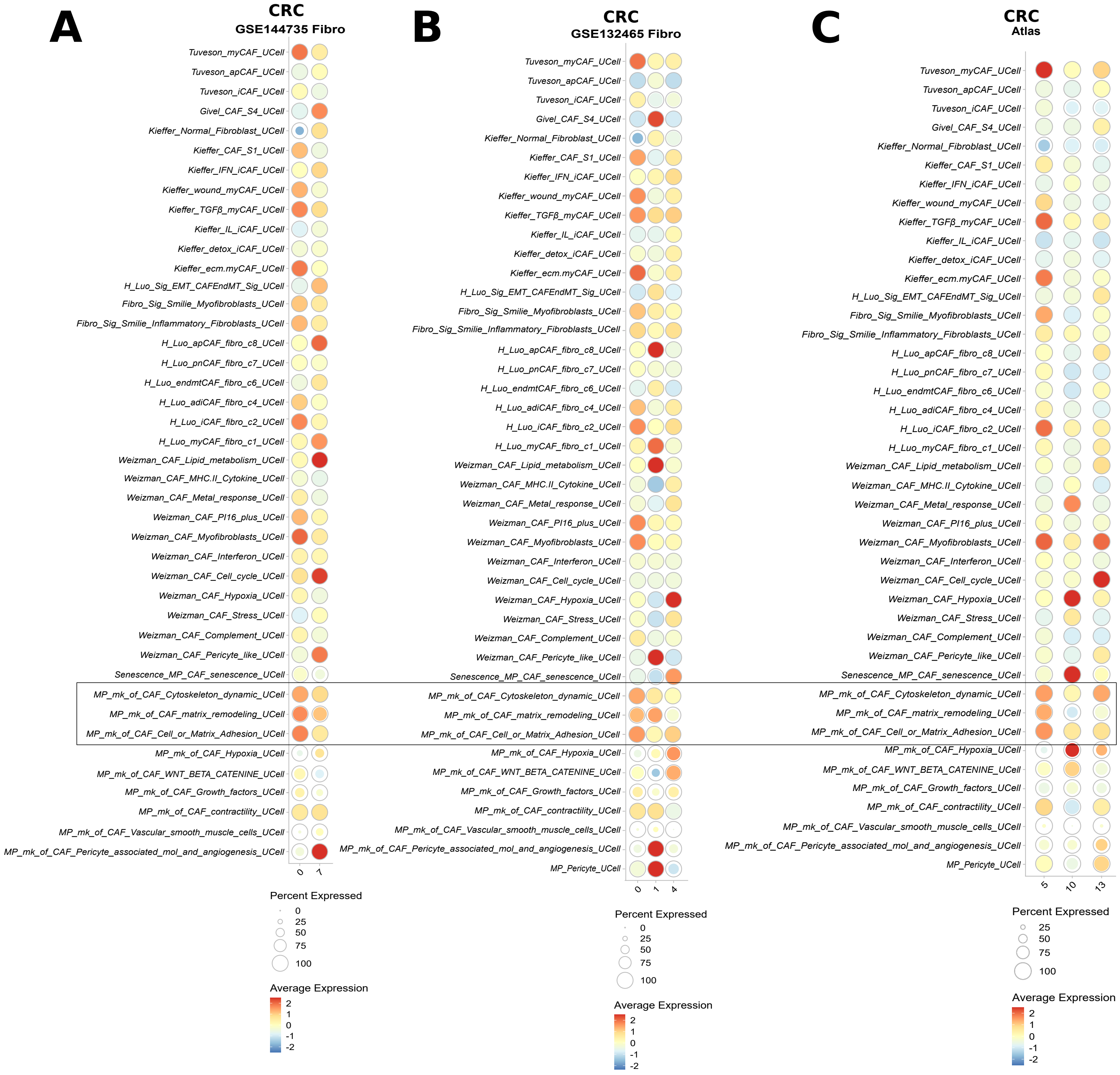

### Supplemental Figure 3

#### Supplementary Figure 3

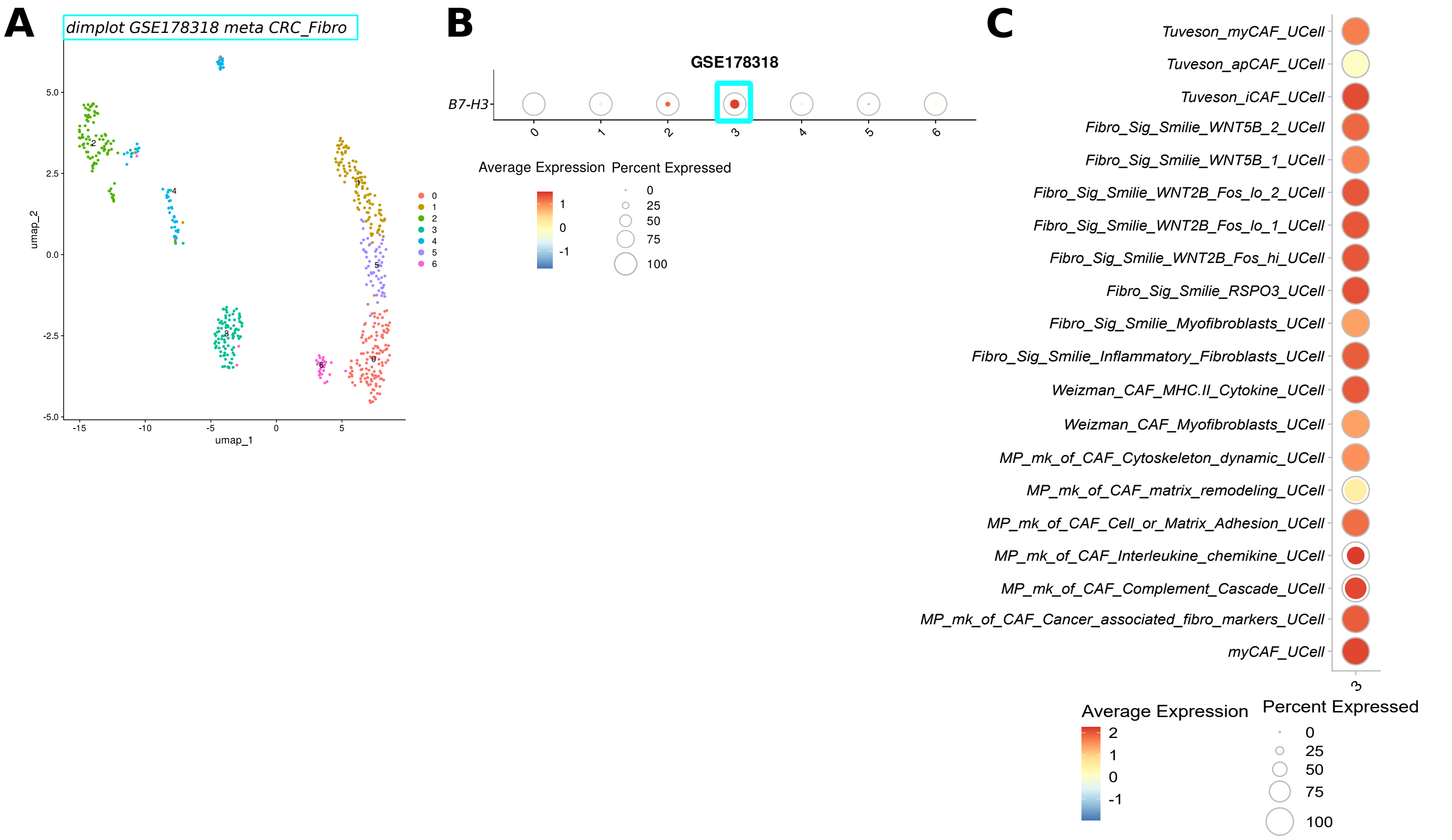
